# Single molecule dynamics at a bacterial replication fork after nutritional downshift

**DOI:** 10.1101/2020.07.29.226316

**Authors:** Rogelio Hernández-Tamayo, Hannah Schmitz, Peter L. Graumann

**Affiliations:** SYNMIKRO, LOEWE Center for Synthetic Microbiology, Marburg, Germany; Department of Chemistry, Philipps Universität Marburg, Marburg, Germany

**Keywords:** replication, single molecule microscopy, DNA primase, helicase, stringent response, *Bacillus subtilis*

## Abstract

Replication forks must respond to changes in nutrient conditions, especially in bacterial cells. By investigating the single molecule dynamics of replicative helicase DnaC, DNA primase DnaG, and of lagging strand polymerase DnaE in the model bacterium *Bacillus subtilis* in response to transient replication blocks due to DNA damage, to inhibition of the replicative polymerase, or to downshift of serine availability, we show that proteins react differentially to the stress conditions. DnaG appears to be recruited to the forks by a diffusion and capture mechanism, becomes more statically associated after arrest of polymerase PolC, but binds much less often after fork blocks due to DNA damage or to nutritional downshift. These results indicate that binding of the alarmone ppGpp due to the stringent response prevents DnaG from binding to forks rather than blocking bound primase. Dissimilar behaviour of DnaG and of DnaE suggest that both proteins are recruited independently to the forks, rather than jointly. Turnover of all three proteins was increased during replication block after nutritional downshift, different from the situation due to DNA damage or polymerase inhibition, showing high plasticity of forks in response to different stress conditions. Forks persisted during all stress conditions, apparently ensuring rapid return to replication extension.

## INTRODUCTION

DNA replication is a highly choreographed and tightly regulated event in the lifecycle of all cells. It is carried out by a dynamic, multi-protein complex known as the replisome (1), which precisely coordinates the action of several distinct factors to efficiently and rapidly couple DNA unwinding with high-fidelity nucleic acid synthesis (2–4). Importantly, DNA replication must respond to situations of changing environmental or developmental conditions, including response to damage in the template, and reduced nutritional capacity of cells (5). This is especially relevant for bacterial cells that are directly affected by changes in their surroundings (6–8).

DNA replication is inherently asymmetric; one daughter strand, termed the leading strand, is continually synthesized in the same direction as the unwinding of the DNA duplex. The other (lagging) strand is synthesized in the opposite direction in short intervals, giving rise to Okazaki fragments (9). The Gram-positive model organism *Bacillus subtilis* and the rest of the Firmicutes (low-G+C-content Gram-positive bacteria) use two essential DNA polymerases, DnaE and PolC, during replication (10,11). *In vivo*, PolC is the main replicative polymerase, while DnaE acts in lagging strand synthesis. *In vitro* reconstitution of the *B. subtilis* replisome has demonstrated that PolC is responsible for all leading strand synthesis as well as most lagging strand synthesis, whereas the more error prone and much slower DNA replicase DnaE (25-60 nt/s for DnaE compared to ~500 nt/s for PolC) plays a crucial role in initiating lagging strand synthesis. DnaE is important for extending the lagging strand RNA primer before handing off to PolC, which then completes replication of the Okazaki fragment (11,12). The synergistic relationship between two polymerases in the *B. subtilis* replisome resembles the synergy found in eukaryotic replication (11,13).

Replication of the lagging strand requires a specialized RNA polymerase, termed primase, to initiate each Okazaki fragment with a short oligoribonucleotide (14). In *B. subtilis* the primase, DnaG, is recruited to the replication fork by an interaction with the replicative helicase, DnaC, where it synthesizes an RNA primer every 1.5–2 kb (1). Working together, the helicase and primase unwind the DNA template and initiate thousands of regularly spaced Okazaki fragments to promote fork progression at a rate of 1000 bases per second in rapidly dividing bacterial cells.

The direct association of primase and helicase co-regulates their functions. For example, primase increases both the ATPase and helicase activities of DnaC. Similarly, DnaC can modulate the overall activity of DnaG, as well as the length of primers synthesized by primase, and its initiation specificity (15). Each Okazaki fragments is initiated by a short RNA primer by DnaG for extension by DnaE. Primase requires interaction with the helicase to stimulate its RNA polymerase activity (16). In *E. coli* the helicase-primase contact is established through the interaction between the C-terminal domain of primase (17,18) and the N-terminal domains of the helicase (19). In *E. coli*, this interaction is weak and transient, with a K_D_ in the low micromolar range, giving rise to fast on/off kinetics (18). Yet, in *Geobacillus stearothermophilus*, helicase and primase form a stable complex that can be isolated and crystallized (18,20).

Bacteria respond to nutrient downshift by the so-called stringent response. Non-charged tRNAs arise when a shortage of any amino acid occurs, and bind to the A-site on the ribosome, where they are sensed by RelA (21,22). This multifunctional enzyme in turn is activated and converts GTP into ppGpp (guanosine 3’,5’-bispyrophosphate) or pppGpp (guanosine 3’-diphosphate 5’-triphosphate), a second messenger that triggers many events including downregulation of translation, and increasing the synthesis of enzymes for e.g. amino acid synthesis pathways (23). Interestingly, (p)ppGpp also binds to DnaG, whereby DNA replication is greatly slowed down, or even stopped, until nutrient shortage is overcome (24). The stringent response induces arrest of DNA replication in *B. subtilis* and, to a lesser extent, in *E. coli* (25,26). Interference of (p)ppGpp with the activity of DNA primase inhibits replication elongation in a dose-dependent manner and adjusts elongation rate according to the nutritional status of the cell (27,28).

We were interested in finding out more exactly how replication forks are adapted during nutritional downshift inducing the stringent response, and in comparing changes in molecule dynamics occurring in response to DNA damage-induced blocks of replication forks, or by specifically blocking PolC. By generating functional fluorescent protein fusions to DnaC and to DnaG, and employing a partially characterized fluorescent protein fusion to DnaE, we find distinct changes at the single molecule level to the three scenarios described above, showing that replication forks have a high plasticity to deal with different stress situations in a bacterial cell.

## MATERIALS AND METHODS

### Bacterial Strains and Growth Conditions

The bacterial strains and plasmids used in this study are listed in Table S1, and the nucleotides are listed in Table S2. *Escherichia coli* strain DH5α (Stratagene) was used for the construction and propagation of plasmids. All *Bacillus subtilis* strains were derived from the wild-type strain BG214. Cells were grown in Luria-Bertani (LB) rich medium at 30°C. When needed, antibiotics were added at the following concentrations (in μg/ml): ampicillin, 100; chloramphenicol, 5; spectinomycin, 100; kanamycin, 30. When required, media containing mitomycin C (MMC), 50 ng/ml; 6(p-Hydroxyphenylazo)-uracil (HPUra), 50 μg/ml; or D,L-Serine Hydroxamate (SHX) 5 mg/ml were prepared by adding appropriate volumes of a filter-sterilized solution.

### Construction of strains

DnaE, DnaG and DnaC were visualized as a DnaE-mVenus, DnaG-mVenus and DnaC-mVenus (“mV”), fusion proteins expressed at the original locus. The last 500 bp coding for each gene were integrated into vector pSG1164-mVenus (29), using *Apa*I and *EcoR*I restriction sites, and BG214 cells were transformed with this construct, selecting for Cm resistance (leading to strains Table S1). For colocalization studies, DnaX-CFP was integrated at *amyE* locus by the use of the plasmid pSG1192 and expression was controlled by xylose addition (30). To investigate colocalization of DnaE, DnaG and DnaC, the resulting strains PG3307, PG3322 and PG3323 (see Table S1) were transformed with chromosomal DNA of strains leading to the expression of DnaX-CFP in parallel to DnaE-mV, or to DnaG-mV, or to DnaC-mV.

### Fluorescence Microscopy

For fluorescence microscopy, *B. subtilis* cells were grown in LB at 30°C under shaking conditions until exponential growth. Conventional light microscopy was performed using a Zeiss Observer Z1 (Carl Zeiss) with an oil immersion objective (100 x magnification, 1.45 numerical aperture, alpha Plan-FLUAR, Carl Zeiss) and a CCD camera (Cool SNAP EZ, Photometrics). Data were processed using Metamorph 7.5.5.0 software (Molecular Devices, Sunnyvale, CA, USA).

### Single molecule microscopy and tracking

Cells were spotted on coverslips (25 mm, Menzel) and covered using 1% agarose pads prepared before with fresh S7_50_ minimal medium by sandwiching the agarose between two smaller coverslips (12 mm Marienfeld). All coverslips were cleaned before use by sonication in Hellmanex II solution (1% v/v) for 15 min followed by rinsing in distilled water and a second round of sonication in double distilled water. In contrast to the wide-field illumination used in conventional epifluorescence microscopy, the excitation laser beam used in our setup is directed to underfill the back aperture of the objective lens, generating a concentrated parallel illumination profile at the level of the sample, leading to a strong excitation followed by rapid bleaching of the fluorophores. When only a few unbleached molecules are present, their movement can be tracked. In addition, freshly synthesized and folded fluorophores become visible when the sample is excited again. When an observed molecule is bleached in a single step during the imaging, it is assumed to be a single molecule (4,31). Image acquisition was done continuously during laser excitation with the electron-multiplying CCD (EMCCD) camera iXon Ultra (Andor Technology, Belfast, UK). A total of 2,500 frames were taken per movie, with an exposure time of 20 ms (23 frames per second [fps]). The microscope used in the process was an Olympus IX71, with a ×100 objective (UAPON 100×OTIRF; numerical aperture [NA], 1.49; oil immersion). A 514-nm laser diode was used as excitation source, and the band corresponding to the fluorophore was filtered out. Of note, cells continued to grow after imaging, showing that there is little to no photodamage during imaging, while cells stop growing when exposed to blue light (below 480 nm). Acquired streams were loaded into Fiji ImageJ (32). Automated tracking of single molecules was done using the ImageJ plugin MtrackJ, or u-track 2.1.3 (33).

### Diffusion analysis of single-molecule tracks

Tracking analysis was done with u-track-2.1.3, which was specifically written for Matlab (MathWorks, Natick, MA, USA). Only trajectories consisting of a minimum of 5 frames were considered tracks and included for further analysis. A widely accepted method to analyse the diffusive behaviour of molecules is by using the mean squared displacement (MSD)-versus-time-lag curve (34,35). This provides an estimate of the diffusion coefficient as well as of the kind of motion, e.g., diffusive, sub-diffusive or directed. However, the method requires that within a complete trajectory there be only one type of homogeneous motion and that the trajectory is preferably of infinite length. To distinguish immobile and mobile molecules from each other, we compare the frame-to-frame displacement of all molecules in x and the y directions. Using a Gaussian mixture model (GMM) to fit the probability density distribution function of all frame-to-frame displacements, determine the standard deviations σ_1_ and σ_2_, as well as the percentages F_1_ and F_2_ of the slow and the fast subfractions of molecules, respectively. Finally, the diffusion constants were calculated according 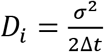, (*i* = 1,2) where *Δt* is the time interval between subsequent imaging frames.

Generation of heat maps, analyses of molecule dwell times, and visualization of slow and fast tracks in a standardized cell are based on a custom-written Matlab script (SMTracker) that is available on request (36). SMTracker can use particle tracking tools u-track (33) and TrackMate (37) and computes the x- and y-coordinates of molecular trajectories relative to the geometry of each cell, as obtained by the cell segmentation tools MicrobeTracker (34) or Oufti (38).

### Dwell times

Dwell time is defined as the average duration that a particle stays inside a certain region. Observing the trajectories in this manner could give insights, for example, on how long the replication proteins are bound at the replication fork. For that matter, dwell time calculations need as parameters the (circular) region, and the minimum number of steps that a molecule should remain inside the region (1 step = 1 interval time). The procedure operates in such a way that searches for the longest dwell events of the protein in each trajectory. For T = (C_1_,…C_n_) a trajectory is defined as a set of nodes Ci = (x_i_,y_i_), and x_i_,y_i_ the nodes’ coordinates in a cartesian axis, the circle C(C_k_,R) is chosen, with R being the radius that contains the max_i_mum number of consecutive points of the trajectory. Then, the amount of time the molecule stays is counted and the same track T excluding that segment of trajectory, T/(C_k_,…C_k+p_), is again searched for more dwell events. The procedure finishes when no more dwell events can be found. In our procedure, one gap (point absent for one frame) or one point outside the circle that goes and comes back are also considered to have remained inside the circle. The number of dwell events and their frequency is plotted in a pdf-histogram, and the data are fitted to a single – or if appropriate, multi-exponential decay, in order to distinguish up to two different populations of dwell times events.

### Statistical data analysis

The goodness of fits of the Gaussian mixture models was assessed using probability-probability plots (pp-plots). Errors on the fitted parameters are given as 95% confidence intervals, which were derived from the Jacobian matrix of the nonlinear optimization process using the MATLAB™ function *nlparci.* To compare fraction sizes and diffusion constants under different conditions and between different proteins, statistical hypothesis-testing was performed using *Z*-tests. Differences in dwell time and step size distributions were tested using a Kolmogorov-Smirnov 2-sample test. To assess the most likely number of populations for each fit, we applied the Bayesian Information Criterion (BIC), as detailed in (36). As usual, ** and *** stands for p-values lower than 0.01 and 0.05 respectively, while n.s. means statistically “not significant”. Statistical hypothesis testing and plotting was performed using SMTracker (36) and MATLAB™ custom written scripts.

## RESULTS

We wished to gain insight into the changes of dynamics occurring at *Bacillus subtilis* replication forks in response to conditions inducing a transient block in DNA replication, including the response to nutritional downshift, termed stringent response. Our strategy was to employ serine hydroxamate (SHX) to induce the stringent response (SHX blocks serine tRNA synthetase, leading to an accumulation of uncharged serine tRNAs) (39) and to compare dynamics of replication proteins to those seen after addition of Mitomycin C (MMC), which induces DNA damage supposed to transiently block the progression of forks (11,40,41), or of 6(p-Hydroxyphenylazo)-uracil (HPUra), which reversibly binds to and inhibits DNA polymerase PolC, thereby blocking progression of replication (42).

We additionally generated functional mVenus fluorescent protein fusions to DnaC and to DnaG, and employed fusions to DnaE and to DnaX, previously shown to be able to functionally replace wild type proteins (43). We found that a fusion of mVenus to the C-terminus of DnaC or to that of DnaG, each expressed from the original gene locus under native transcriptional control, as sole source of the protein in the cell, did not negatively affect exponential growth of *B. subtilis* cells (Fig. S1A and C). Additionally, treatment of cells expressing each fusion with MMC, HPUra or SHX did not compromise viability (Fig. S1E). Note that DnaE-mVenus expressing cells showed a weak sensitivity towards SHX treatment (Fig. S1B and S1E). Using high resolution epifluorescence microscopy, we found that very similar to DnaX-CFP, DnaC-mVenus formed fluorescent foci within the cells (Fig. 1A). In 80% of cases, DnaX-CFP and DnaC-mVenus foci colocalized (in the rest of events, only one of the two signals were visible), showing that the helicase fusion is recruited to replication forks, as expected.

**Fig. 1.**
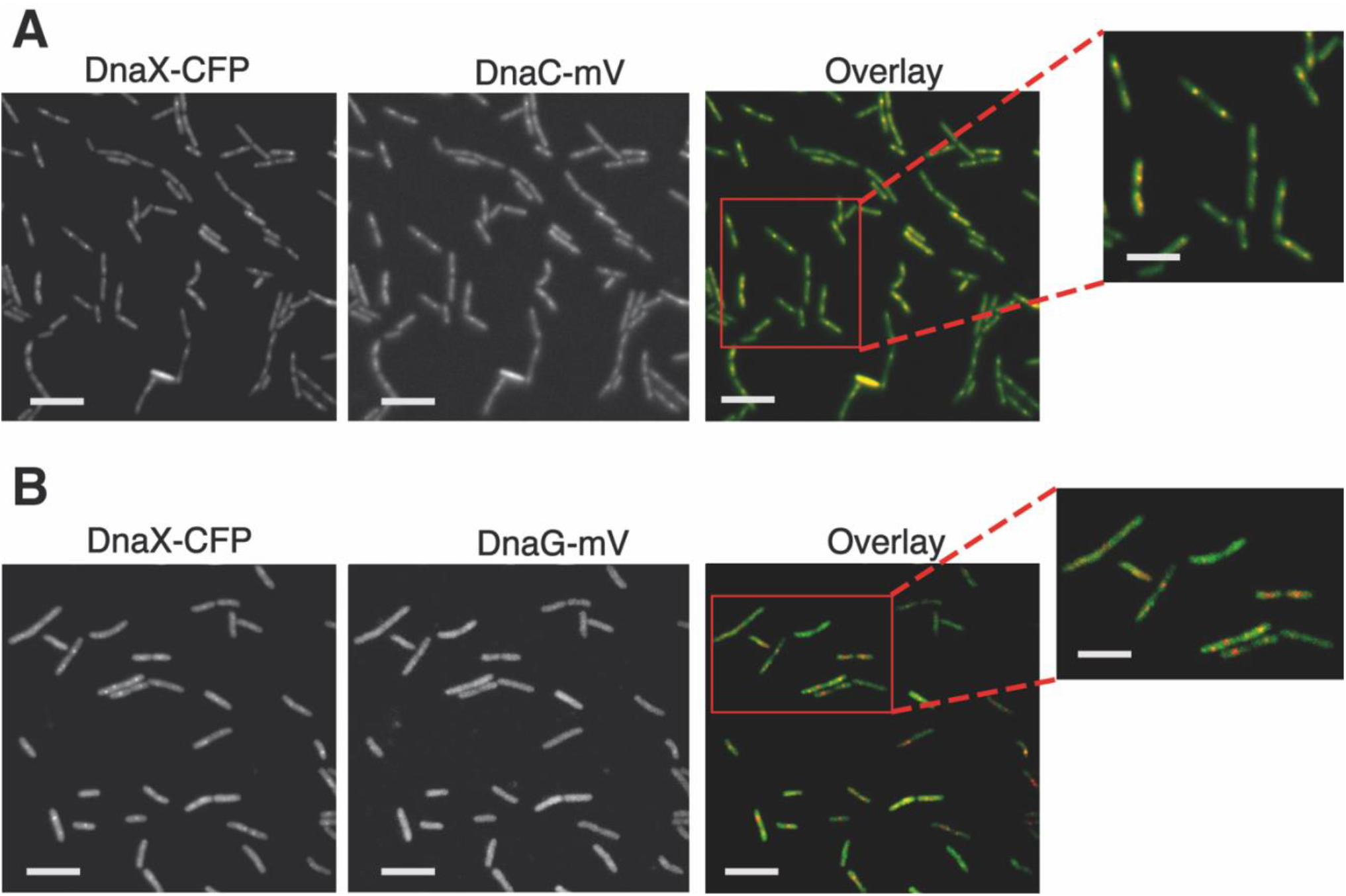
Colocalization of DnaC-mVenus, or of DnaG-mVenus with DnaX-CFP in *B. subtilis* cells. Epifluorescence microscopy of cells expressing A) both DnaC-mV and DnaX-CFP, or B) DnaG-mV and DnaX-CFP, during exponential growth as sole source of the proteins. Scale bars 5 μm and zoom panel 2 μm.

For DnaG, two principle scenarios could be envisioned: a) DnaG may come and go to forks by a diffusion/capture mechanism, which would lead to an exchange event every few thousand base pairs, b) DnaG might have binding sites at the forks (analogous to DnaA, the initiator of replication, (44)), which would lead to an enrichment at the forks and a concomitant enhanced replacement efficiency through a local pool of molecules. Fig. 1B shows that in most cells, DnaG-mVenus was dispersed throughout the cells, with some cells showing weak accumulations. Thus, at first sight, there does not seem to be an accumulation of DnaG at the forks.

Next, we treated cells with concentrations of MMC, HPUra or SHX that led to slowed down growth rate but did not stop growth (Fig. S1A-C). We reasoned that these concentrations were able to strongly act at the respective targets including replication forks but did not lead to a large degree of cell death. Fig. 2 shows that DnaC-mVenus foci appeared to be visually weaker after treatment with HPUra and SHX than after MMC treatment. As we will move on to single molecule tracking (SMT) below, we will refrain from quantifying at this place. For DnaG-mVenus, we noted the appearance of visible foci after addition of MMC and of HPUra, but not in response to SHX (Fig. 2). While there were qualitative changes in localization patterns, we clearly observed the continued presence of foci representing replication forks for the three analysed conditions, indicating that in general, forks persist through the three types of treatments.

**Fig. 2.**
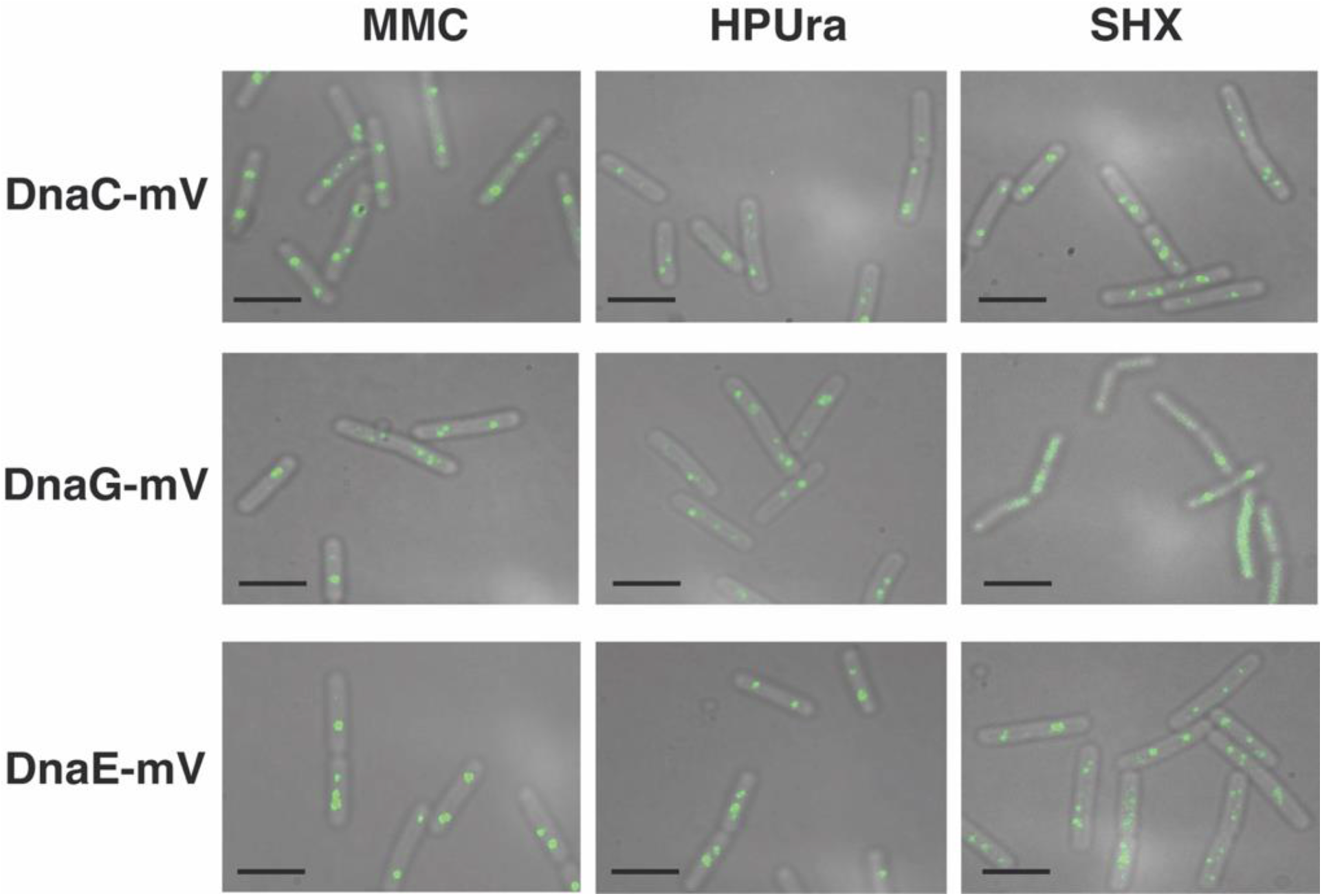
Localization of DnaC-mVenus, DnaG-mVenus and DnaE-mVenus during stress conditions. Epifluorescence microscopy of *B. subtilis* cells expressing DnaC-mV, DnaG-mV or DnaE-mV as sole source of the proteins treated with different drugs as indicated above the panel rows. Scale bars 2 μm.

### Single molecule tracking reveals poor enrichment of DnaG at replication forks, arguing for a diffusion capture mechanism for recruitment

We next turned towards SMT to quantify changes in protein dynamics at a single molecule level, using an experimental setup that has been described before (43,45,46). We used SMTracker software 1.5 (36) to analyse tracks that were generated using u-track software (33). We employed 20 ms stream acquisition to ensure that even freely diffusing molecules of DnaC (50.4 kDa plus mVenus) and of DnaG (68.6 kDa plus mVenus) would be trackable. Fig. 3A and 3B show overlays of all frames of a typical movie (see movies S1 and S2 for examples) of DnaC, revealing clear foci within cells that contain little background, indicative of a high number of DnaC molecules being present at the forks. Conversely, we observed barely detectable discrete accumulations of DnaG, but many molecules localized throughout the cells. Thus, these two proteins show clearly distinct localization patterns, indicating that DnaG only transiently and shortly associates with the replication machinery, contrarily to DnaC. For quantification of molecule trajectories, we employed Gaussian Mixture Modelling (GMM), which allows to directly compare molecule dynamics between different proteins, or for a given protein between different growth conditions: a common value for the diffusion constant “D” is found, which leaves alterations only to occur in changes of the population size of molecules with a given diffusion coefficient. GMM also allows to distinguish if the probability density function of observed step sizes can be explained by a single Gaussian function, and thus by the presence of a single population of molecules having the same value for D, or by two or three different populations, which is tested by an r2 analysis (47). Fig. S2 shows that for all three proteins, observed distributions could be well explained with the existence of two populations, but not of one. One fraction had a low diffusion constant, corresponding to molecules bound to replication forks, and one with a high value for D, characteristic of freely diffusing molecules, as was described before (43). Fig. 4, upper panel illustrates that DnaC featured the highest proportion of statically bound molecules, with 81% (Fig. S3), and only 19% diffusive molecules, as was expected for the helicase. DnaE showed a relatively balanced proportion of 58% for bound and 42% diffusive molecules (Fig. S3), while DnaG had the lowest population of static molecules (46%), and a larger amount (54%) of unbound ones. These data strongly suggest that DnaG comes and goes to forks, and is only present at very low, likely single molecule level at the replication machinery, suggesting a diffusion capture model for recruitment to the lagging strands, rather than an exchange with molecules pre-bound to other replication-associated proteins.

**Fig. 3.**
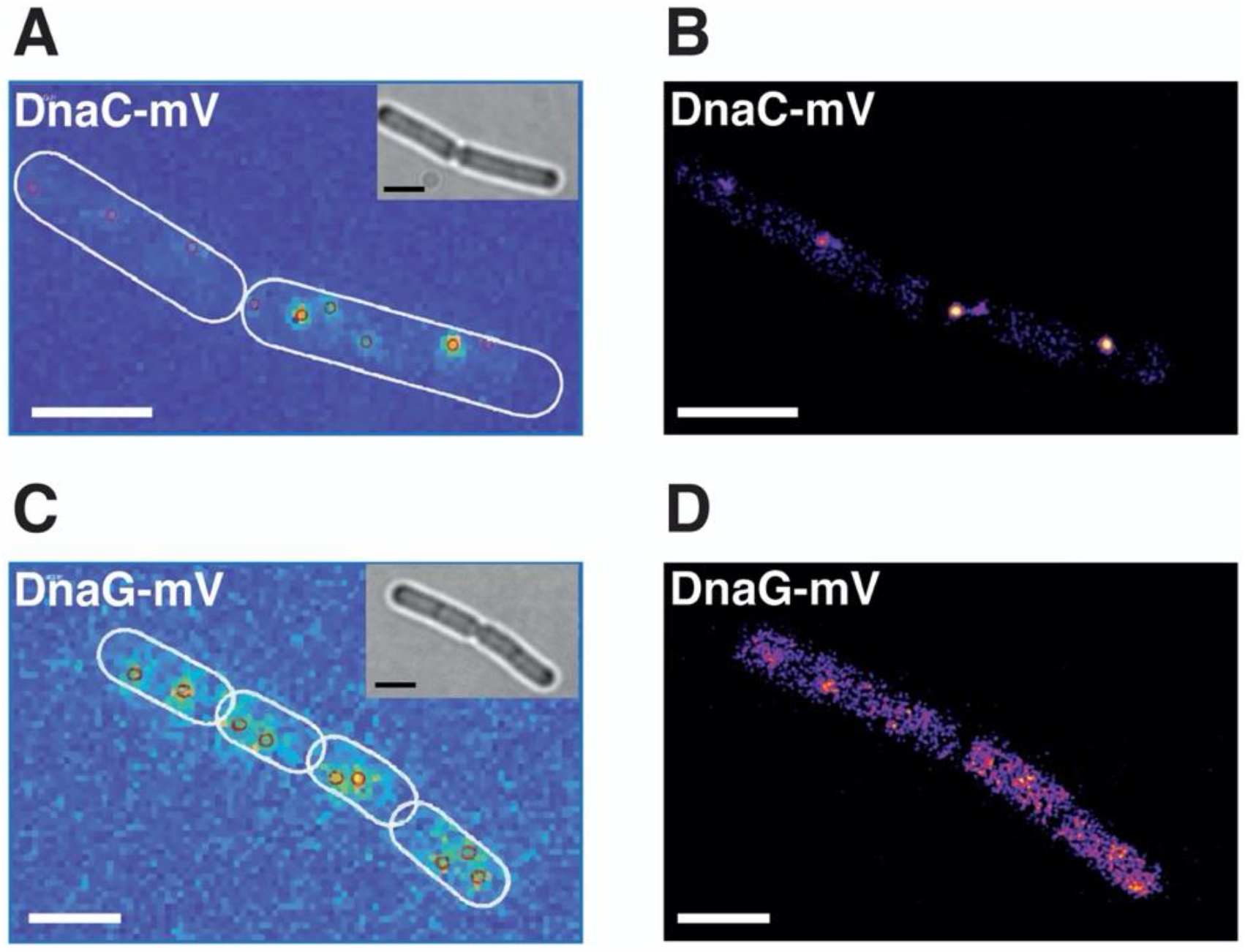
Sum of single molecule tracking movies of exponentially growing cells expressing DnaC-mVenus or DnaG-mVenus. Insets in panels A and C show bright field images, outlines of cells are indicated by white ovals. Panels A and C show heat maps of localization, panels B and D fluorescence images. White bars 2 μm.

**Fig. 4.**
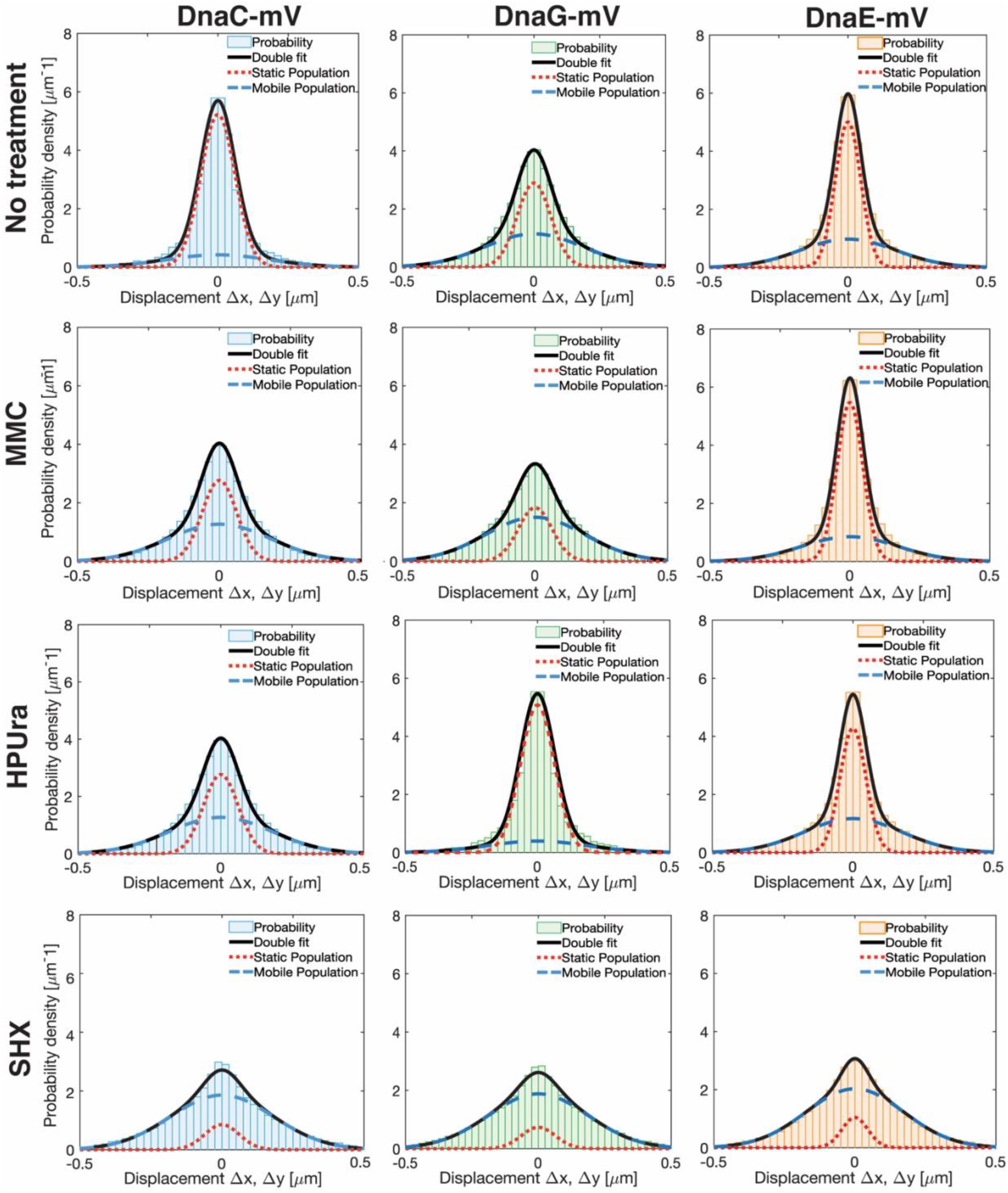
Diffusion patterns of DnaE-mVenus, DnaC-mVenus and DnaG-mVenus. Gaussian mixture model (GMM) analyses of frame-to-frame displacements in x- and y-directions. Black lines represent the sum of the two Gaussian distributions. Dotted red and blue lines represent the single Gaussian distributions corresponding to the static and mobile fractions.

### DnaC, DnaG and DnaE respond differentially to stress conditions at a single molecule level

Addition of MMC to growing cells did not strongly affect the size of DnaE populations, as was described before (43). If anything, DnaE turned to be slightly more stably associated with the forks, becoming more static from about 58 to 63% (Fig. S3). Conversely, both DnaC and DnaG revealed a decrease in the static population with a concomitant increase in unbound molecules (Fig. 4, second row of panels). For DnaC, the size of the static population observed during exponential growth (about 81%) was almost halved to 42%, while for DnaG, the static population decreased from ~ 46 to 29% (Fig. S3). Because chromosome segregation and thus replication continue after addition of MMC, albeit more slowly (48), likely based on frequent restart processes, it follows that DnaC molecules are more frequently exchanged during replication restart than polymerases. Unexpectedly, blocking of PolC by HPUra lead to an increase of fork-bound primase molecules (Fig. 4, third row of panels), while populations of DnaC remained unchanged, and DnaE became slightly more dynamic. The static fraction of DnaG almost doubled from ~ 46 to 82%, while that of DnaC almost halved from ~ 81 to 42% (Fig. S3). Thus, DnaC and DnaG respond in an opposite manner to blocking of polymerase activity. Interestingly, under this condition, DnaE was partially lost from forks, with the static fraction dropping from ~ 58 to 49% (Fig. S3).

Induction of the stringent response induced a third pattern of changes, namely a strong decrease in fork-bound, static molecules and thus a large increase in freely diffusive molecules for all three replication proteins. We interpret these findings to mean that a) stringently blocked forks become highly prone to protein exchange (in agreement with weaker fluorescence observed using epifluorescence microscopy, Fig. 2, b)) in spite of a block in DnaG recruitment, forks do not completely disintegrate, but are likely ready to rapidly return to activity when ppGpp levels are lowered.

The different behaviour of DnaG and DnaE at a single molecule level after the different kinds of replication stress induced suggests that they are independently recruited to replication forks. Were they generally recruited in an ensemble manner, we would have expected similar shifts in bound and free populations of molecules during the same stress conditions.

### Nutritional downshift increases turnover of helicase, primase and polymerase at the forks

We wished to obtain more information on the exchange rates of the three replication proteins under investigation at the replication machinery. We therefore introduced into SMTracker an analytical tool to quantify the extent of molecules that show transitions between mobile mode and static behaviour. Confined motion was defined as molecules staying within a radius of 120 nm (about three times our localization error) for at least 9 steps (Fig. 5A). These states of static motion are indicated in red colour in Fig. 5. Note that a confined track can be part of a longer track that changes between static and mobile mode, or vice versa (Fig. 5A). Such “transition” tracks are shown in green in Fig. 5. Not shown are tracks that are entirely mobile, and do not rest for 9 steps in a row (see Fig. S4); note that even freely diffusive molecules can stochastically stop for shorter periods of time. Conversely, confined motion occurs when a protein has restricted movement for an extended amount of time, which is due to an interaction/binding to a much larger subcellular structure. To locate these events, a confinement map tool was developed, using the information given by the dwell time calculation and projecting events into a standardized cell. A trajectory is considered to present confinement when it has at least one dwell event (confined track), transitions (mixed behaviour) are changes between confinement and mobility, freely diffusive track lack any kind of confinement. Fig. 5C shows that confined motion of DnaC, DnaG and DnaE clusters in the cell centre where replication forks are present, as expected. In agreement with the finding that DnaG shows the smallest static fraction, it also showed the smallest extent of confined motion, while DnaE showed the largest (Fig. 5C). Treatment with MMC visually increased the percentage of molecules showing transitions, while HPUra did not appear to strongly affect the ratio between confined tracks and those undergoing transitions (Fig. 5C). To better quantify these findings, we scored tracks undergoing purely diffusive motion, transitions or purely confined motion. The latter two fractions were set to 100%, and we scored the change between confined and transitory molecules. Fig. 5B shows that MMC treatment slightly increased transitions of DnaC at the forks, this was more pronounced during blocking of polymerases by HPUra, in agreement with DnaC becoming more mobile under this condition (Fig. 4 and S3). Most notably, transitions greatly increased after addition of SHX, not only for DnaC, but also for DnaG and for DnaE (Fig. 5B and C), revealing that during stringent response, all three proteins revealed highly increased exchange rates at the forks, even though these were blocked for elongation. Intriguingly, the pattern of localization of confined and transitory tracks changed markedly under stringent conditions. All three proteins showed localization away from the cell centre towards the periphery of cells (Fig. 5). Curious about this finding, we stained cells with DAPI to study the subcellular localization of the chromosome(s). Fig. S5 shows regular nucleoids during exponential growth, and condensed nucleoids following addition of MMC, as has been reported before (49). Nucleoids were also more condensed after addition of HPUra but were almost completely decondensed during the stringent response (Fig. S5). Previously, it has been shown that 70S ribosomes occupy nucleoid-free zones at the cell poles and at the cell periphery (50), which is lost upon inhibition of transcription, where the nucleoids decondense (51). Our finding that the stringent response also leads to a strong nucleoid de-condensation suggests that also under this condition, the separation between RNA polymerase and ribosomes is lost. Our data also suggest that the central positioning of replication forks is lost, which agrees with the appearance of more randomly located forks seen in epifluorescence (Fig. 2). Thus, nutritional downshift not only strongly increases exchange of proteins at the replication forks, but also affects their subcellular localization.

**Fig. 5.**
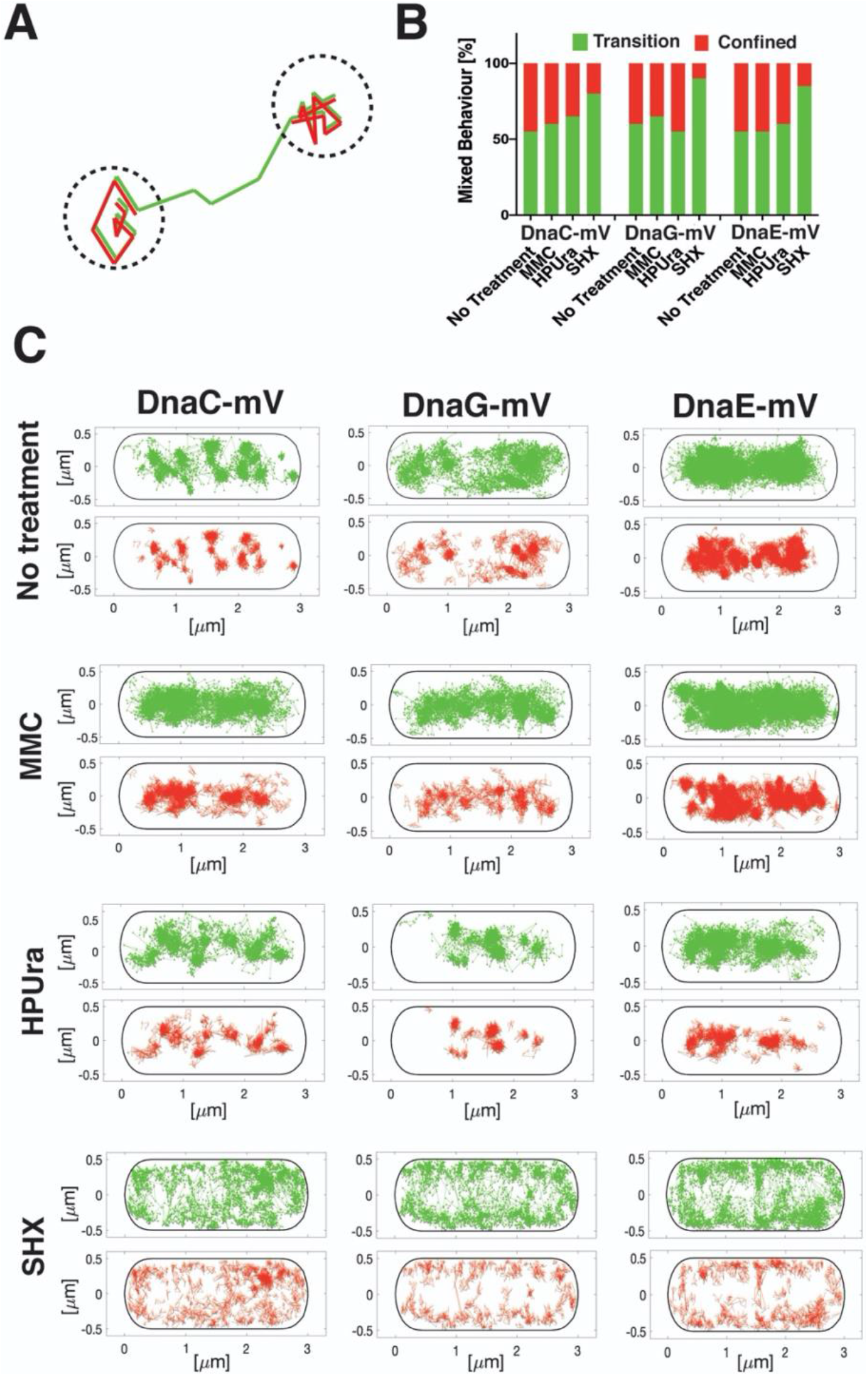
Analyses of tracks for the percentage of transitions under stress conditions. A) cartoon showing the mode of analyses of confined movement (red) versus transition events (green). B) Bar plot showing changes between confined and transitory motion for the three proteins under different stress conditions. C) Maps showing intracellular location of confined motion (as defined by not leaving a radius of 120 nm for at least 9 steps) and transition events for DnaC-mVenus, DnaG-mVenus and DnaE-mVenus, projected into a standardized cell of 3 x 1 μm size.

### Stress conditions differentially affect dwell times of DnaC, DnaG and DnaE

Based on the above findings, we would have predicted lower dwell times of replication at the forks during nutritional downshift. We analysed times molecules dwell within a radius of 120 nm and used a two-population fit to analyse decay curves (Fig. S6). Fraction τ_1_ shows shorter average dwell times and will mostly consist of mobile molecules that stochastically stay put for a short time. Fraction τ_2_ will likely by composed of molecules residing at the replication machinery. Of note, although average track lengths (around 8 steps) were shorter than average dwell times determined, especially for the second fraction (τ_2_) that shows long dwell times, there were enough tracks longer than average track length to allow for a correct extrapolation of average residence time. Because we used YFP-bleaching type SMT, our determined numbers are underestimates of actual dwell times *in vivo*. However, for the sake of comparison between different growth conditions, our estimates are useful to observe relative changes in dwell times. Fig. 6 shows that in agreement with our expectations, the dwell time of DnaC decreased during (transient) replication arrest due to MMC or HPUra treatment, and further decreased during the stringent response. For DnaG, we observed an increased residence time after addition of HPUra (Fig. 6) (Table S3), corresponding to the increase in the fraction of static molecules observed (Fig. 4). However, during stringent conditions, average residence time remained similar to values during unperturbed growth, which is unexpected. DnaE showed clearest changes in dwell times, which decreased from MMC to SHX treatment, where residence times were shortest.

**Fig. 6.**
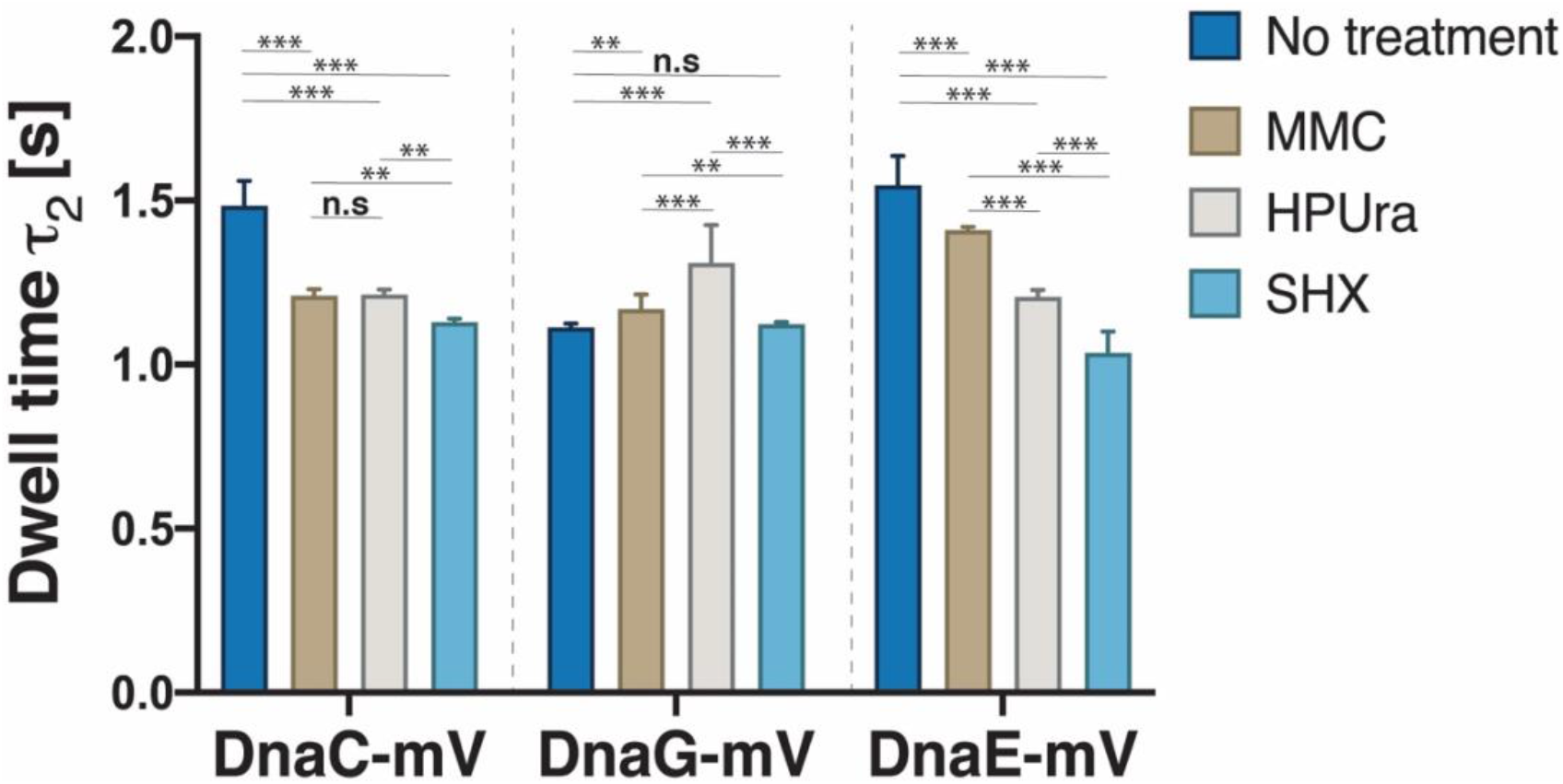
Dwell times. Cumulative distribution of residence times of DnaC-mVenus, DnaG-mVenus and DnaE-mVenus strains, before and after treatment with MMC, HPUra or SHX. Dwell times are estimated using an exponential decay model. Histograms show events of resting fitted by a two-component exponential function. Bars represent long dwell times of DNA-bound molecules. Dark blue bars untreated cells, brown MMC-treated cells, grey HPUra-treated cells and blue light SHX-treated cells. As usual, ** and *** stands for p-values lower than 0.01 and 0.05 respectively, while *n.s.* stands for statistically not significant.

**TABLE 1.**
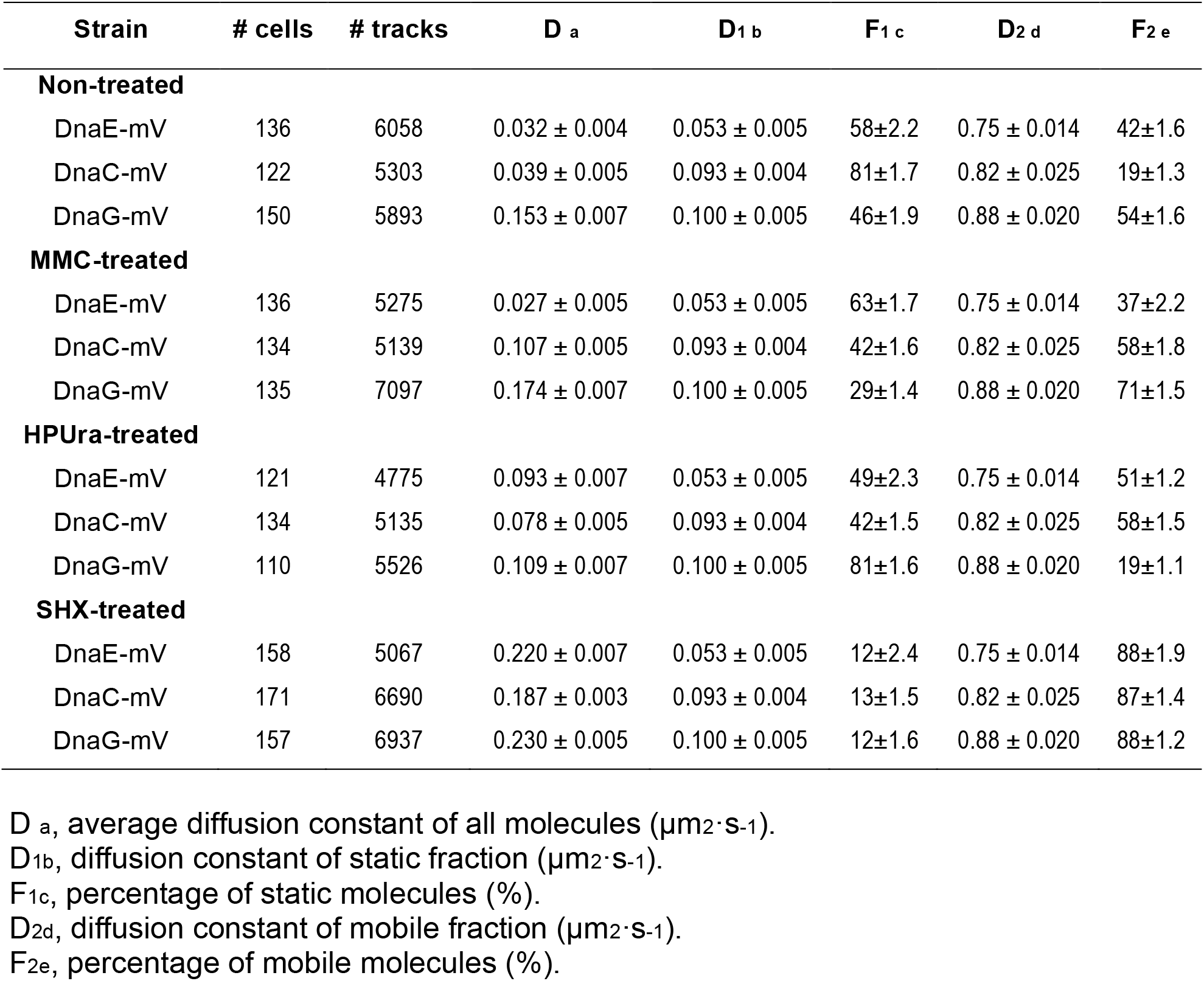
Diffusion constants and percentages of static and mobile molecule fractions.

Keeping in mind the caveat that our determined dwell times are underestimates, we can still observe that DnaC and DnaE show similar residence times during non-perturbed growth. Thus, as DnaE exchanges with every 1000 to 2000 base pairs synthesised at the lagging strand, so do DnaC subunits appear to exchange at this DNA strand. We propose that this exchange is based on exchange of subunits from the hexameric helicase, which thus appear to exchange within a time frame of few seconds.

## DISCUSSION

All cells need to adjust their decision when to replicate to the nutritional state of the cell, as well as to many other conditions and physiological requirements. Especially for bacteria, it is not only important to regulate initiation of replication, but also extension, because a run-out of e.g. nucleotides might be detrimental if replication fork speed was not down regulated. We have sought to shed light onto the question how *Bacillus subtilis*, a model organism especially for the large group of Gram-positive bacteria, adjusts replication at a single molecule level, in response to amino acid starvation. Interestingly, the architecture of *B. subtilis* replication forks is rather similar to that of eukaryotic cells, and dissimilar to other model bacteria such as *E. coli*, in that two replicative polymerases act at the lagging strand (43,52–54), besides other differences.

Recent studies of bacterial DNA replication have led to a picture of the replisome as an entity that freely exchanges DNA polymerases and displays intermittent coupling between the helicase and polymerase(s) (9). Challenging the textbook model of the polymerase holoenzyme acting as a stable complex coordinating the replisome, these observations suggest a role of the helicase as the central organizing hub. We show here that there is a high degree of plasticity in the interaction between the lagging strand polymerase and the replicative helicase upon association of the primase with the replisome. By combining epifluorescence and *in vivo* single-molecule assays, we demonstrate that replicative helicase DnaC, DNA primase DnaG, and lagging strand polymerase DnaE act differentially in response to transient replication blocks due to DNA damage, to inhibition of the replicative polymerase, or to downshift of serine availability (stringent response).

Addition of HPUra has been shown to block the activity of the replicative DNA polymerase in several bacterial species. Interestingly, we find more static binding of primase DnaG at the forks, and an increase in its dwell times, while DnaE becomes slightly more dynamic, and helicase DnaC even more so. These findings suggest that blocking of PolC allows for completion of an Okazaki fragment and permits DnaG to reinitiate binding, but slows down or blocks its turnover. Exchange of DnaC molecules was found during exponential growth and in an increased manner during all three stress conditions, when the progression of the forks was blocked or strongly reduced. We interpret these findings to indicate that the hexameric helicase exchanges its subunits within few second intervals, replacing them continuously, as is known from exchange of e.g. rotor parts of the bacterial flagellum (55).

While DnaE became slightly more statically associated with replication forks during MMC-induced DNA repair and less so during blocking of PolC via addition of HPUra, DnaG became highly stabilized during block of PolC activity, and less statically positioned during DNA repair. Thus, responses in single molecule dynamics were quantitatively different between DnaG and DnaE during conditions of replication stress. Likewise, dwell times changed in very different manners during stress conditions, indicating that both proteins are recruited separately and independently from each other to initiate new Okazaki fragments. Moreover, although DnaC, DnaG and DnaE have been shown to form a stable complex *in vitro* (53), changes in single molecule dynamics of DnaC in response to MMC or HPUra treatment were distinct from those of DnaG and DnaE, showing intriguing plasticity in protein dynamics within this complex, and thus within replication forks.

Most pronounced changes in single molecule dynamics were found after nutritional downshift. All three proteins became much less statically associated with forks during the stringent response, concomitant with a decrease in dwell times for DnaC and for DnaE, and strongly increased turnover of binding/unbinding events. These findings show that interaction of DnaG with the stringent response second messenger (p)ppGpp, which is synthesized via RelA in response to binding of uncharged tRNAs at the ribosome A site, strongly reduces binding of DnaG to the lagging strand, rather than blocking its activity and stalling DnaG at the forks. Additionally, our findings show that while chromosomes decondense during the stringent response, in contrast to condensation during fork block via MMC or HPUra, replication forks persist, but feature increased protein turnover. This will lead to slowing down of replication elongation, or even halt replication completely, but allow for rapid regaining of extension, possibly without the need for restart.

We have recently shown that chromosome segregation and thus also DNA replication, which occur concomitantly, are relatively robust against DNA damage-induction via MMC or inhibition of DNA gyrase, continuing to follow a general pattern that resembles that of directed diffusion (48). Here, we show that replication forks can be seen to persist during MMC treatment, as well as during inhibition of PolC or of DnaG via (p)ppGpp binding. Apparently, while featuring rapid exchange of subunits, the replication machinery appears to be very robust against different kinds of stresses and can be tuned down in speed after nutritional downshift. Clearly, nature has evolved a highly adaptable and overall processive/stable machinery for one of the most central aspects of life, duplication of the genetic information.

## ACKNOWLEDGMENTS

This work was supported by the Center for Synthetic Microbiology, SYNMIKRO, at the Philipps-Universität Marburg, by the research consortium MOSLA, funded by the LOEWE Program of the state of Hessen, and by Deutsche Forschungsgemeinschaft (DFG)-funded research consortium TRR 174.

## AUTHOR CONTRIBUTIONS

R. H.-T. conceived of the study, performed all experiments, evaluated data and co-wrote the manuscript. H.S. contributed to experiments shown in Fig. 4 and in Fig. S3 and helped writing the manuscript. P.L.G. conceived of the study, helped to evaluate data and co-wrote the manuscript.

