## Supplementary Material for "Single molecule dynamics at a bacterial replication fork after nutritional downshift"

### Supplementary movies

Movie S1 Stream acquisition (20 ms intervals) of cells expressing DnaC-mVenus as sole source of the protein during exponential growth, images are shown at 50 frames/s (real time).

Movie S2 Stream acquisition (20 ms intervals) of cells expressing DnaG-mVenus as sole source of the protein during exponential growth, images are shown at 50 frames/s (real time).

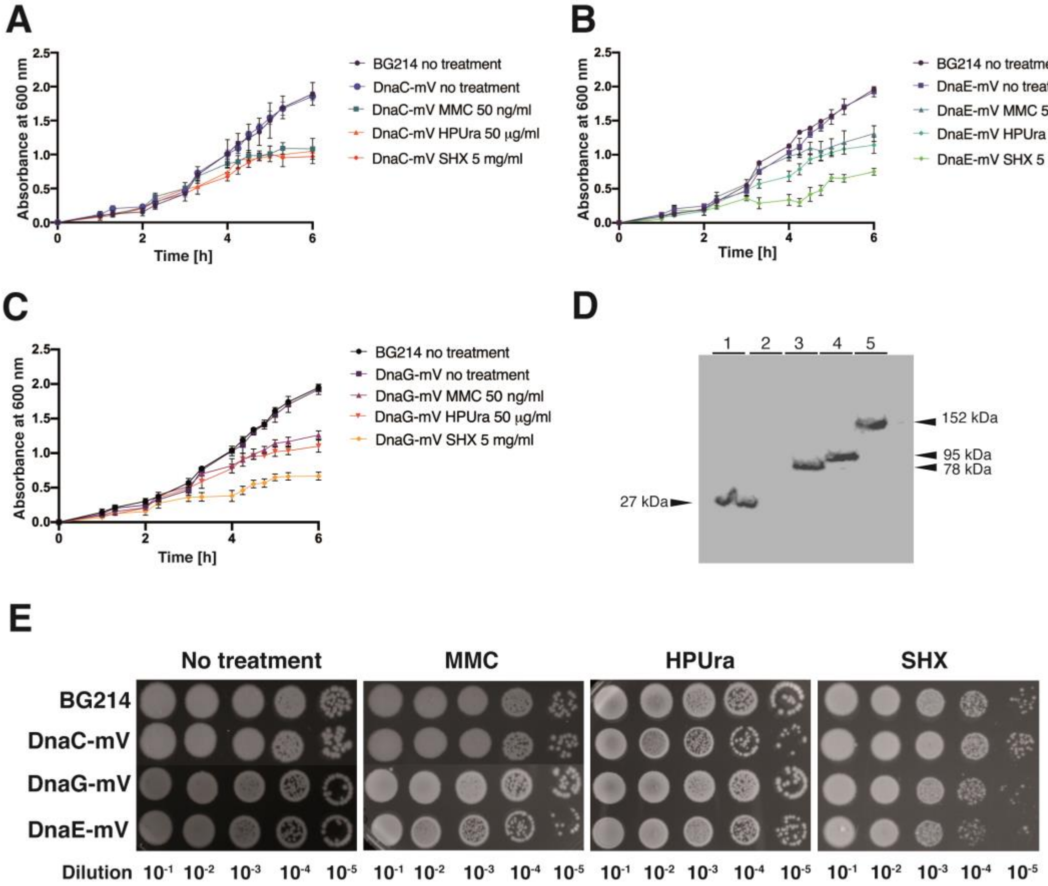

**Fig. S1 A-C)** Survival assay. Cultures of BG214, DnaC-mVenus, DnaE-mVenus and DnaG-mVenus strains, before and after treatment with MMC, HPUra or SHX, with concentrations as indicated in the corresponding legend to the panel. All strains were plated in duplicate, in three independent experiments. **D)** Western blot of fluorescent protein fusions. Shown are Western blots from whole cell lysates of 1) *E. coli* expressing mVenus, 2) *B. subtilis* BG214, or 3) DnaC-mVenus, 4) DnaE-mVenus and 5) DnaG-mVenus expressing cells. Corresponding protein sizes are indicated by arrowheads. Proteins were probed using a 1:500 dilution (rabbit-α-GFP) and secondary antibody was added (goat-α-rabbit-antibody in 1:10000 dilution) after a series of washing steps. Cells were harvested in exponential phase at $OD_{600}$ 0.5 – 0.7 prior to analysis. **E)** Spot assays. Cultures of BG214, or of strains expressing DnaC-mVenus, DnaE-mVenus or DnaG-mVenus, before and after treatment with MMC 50 ng/ml, HPUra 50 µg/ml or SHX 5 mg/ml.

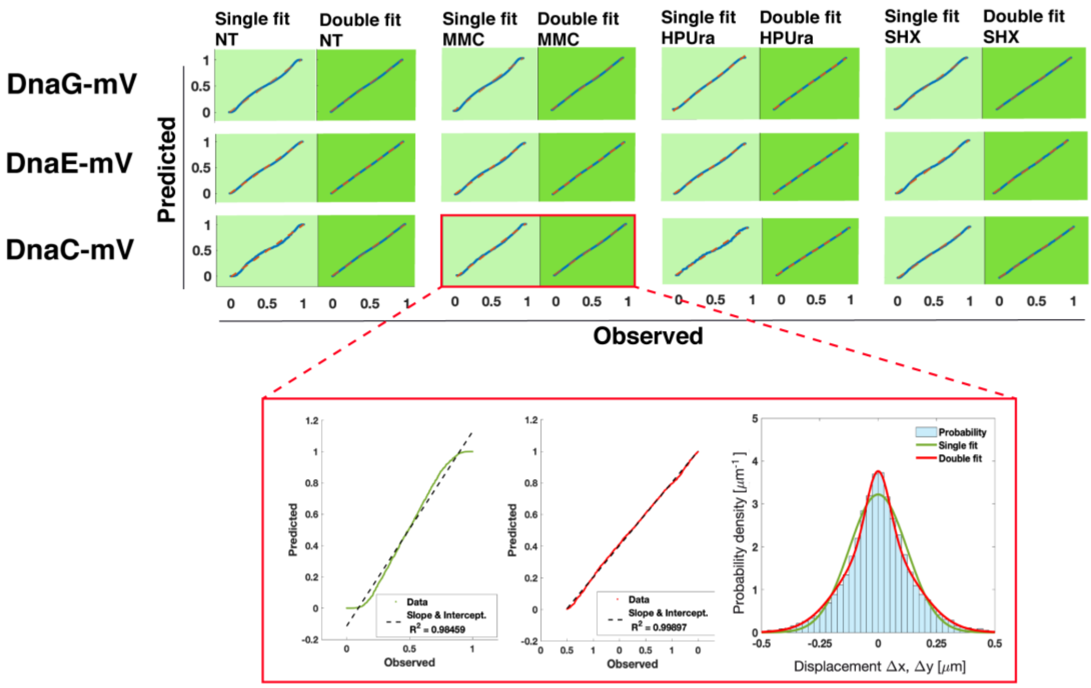

**Fig. S2 Goodness of GMM fit and best model selection.** Probability-probability plots for every type of damage and for every fluorescent protein fusion are displayed. Dark green shows the model that performs better than the other one, in light green the poorer fitting model. Mean Squared Error (MSE) and R-squared were used to find that a two-population fit clearly performs better for all cases. On the highlighted area, details of the correlation and step-size histogram are shown along with the fit for both models, below the panel. A three populations model was discarded for reasons of overfitting. R-squared coefficients were higher than 0.998 for every two-populations model.

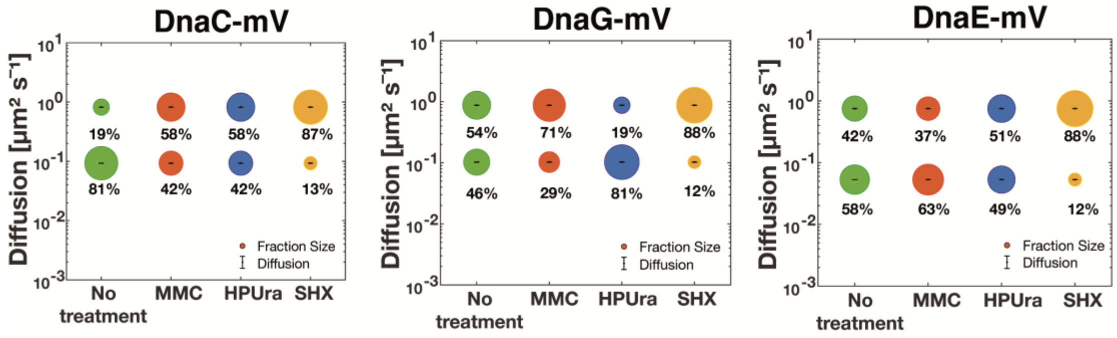

**Fig. S3 Diffusion patterns of DnaC-mVenus, DnaG-mVenus and DnaE-mVenus.** Gaussian mixture model (GMM) analyses of frame-to-frame displacements in x- and y-directions. Bubble plots show a comparison of fraction sizes (size of the bubble) and diffusion constants (y-axis), between different growth conditions: distribution in untreated cells (green circles), in MMC-treated (red circles) HPUra-treated (blue circles) and in SHX-treated (yellow circles) cells. Step size distributions reveal two populations for each protein, a mobile (upper circles) and a static (lower circles) fraction.

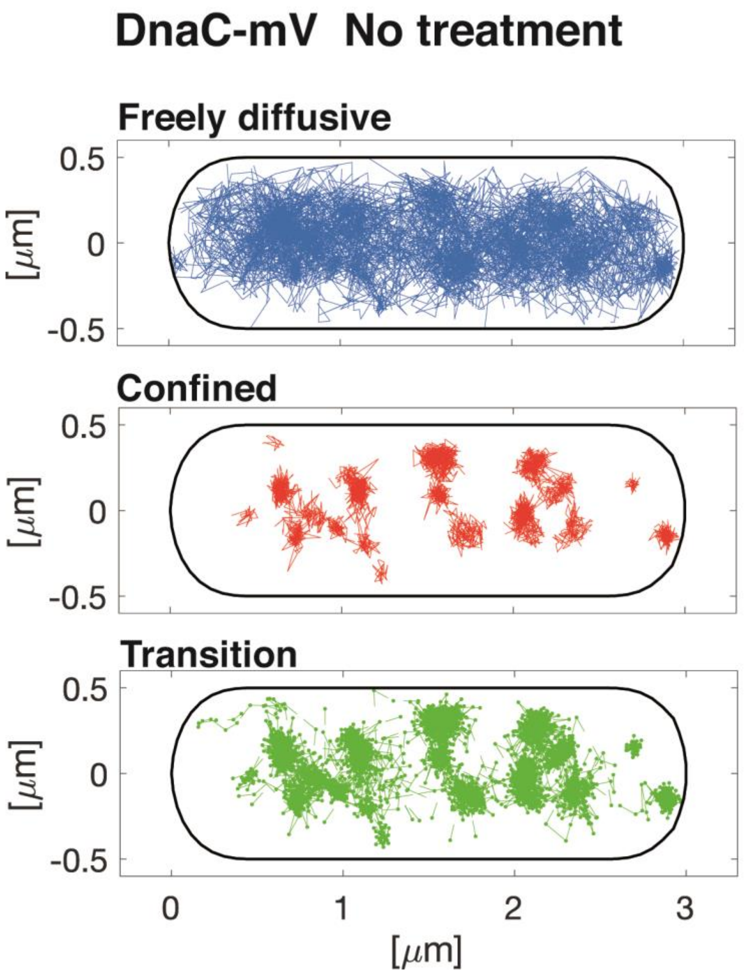

**Fig. S4 Freely diffusive movement of molecules occurs throughout the cell.** A confinement map has been developed alongside the information given by the dwell times calculation algorithm of SMTracker. A trajectory is considered to present confinement (red) when it has at least one dwell event. Molecules changing between confinement and mobility are termed "transition" (mixed behaviour), and is shown in green. Freely diffusive molecules lacking considerable parts of confinement are shown in blue.

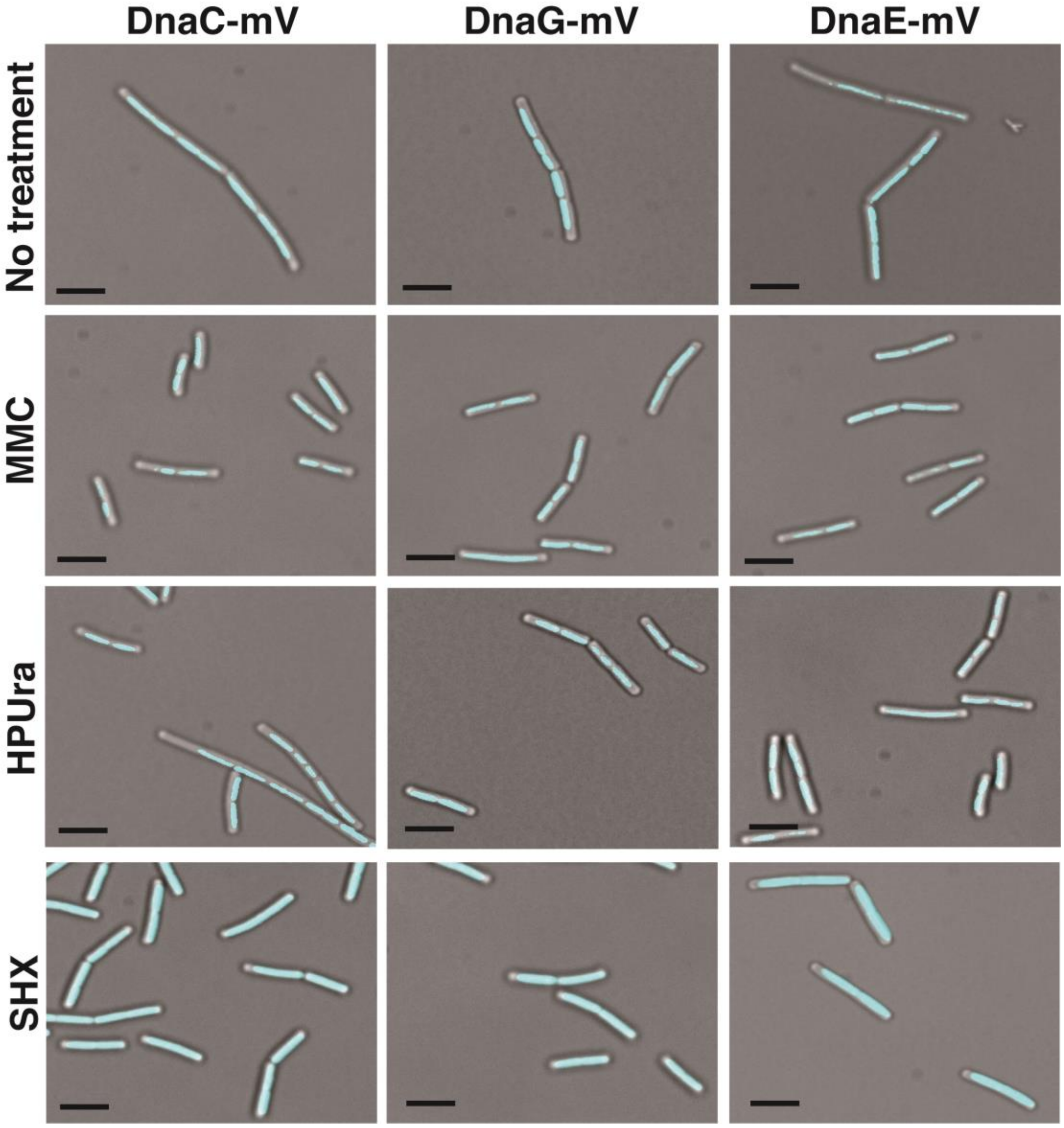

**Fig. S5 DNA localization patterns**. Localization by epifluorescence in representative live *B. subtilis* cells in untreated cells, in MMC-treated, HPUra-treated or in SHX-treated cells as stated next to the panels. Black scale bars 2 µm.

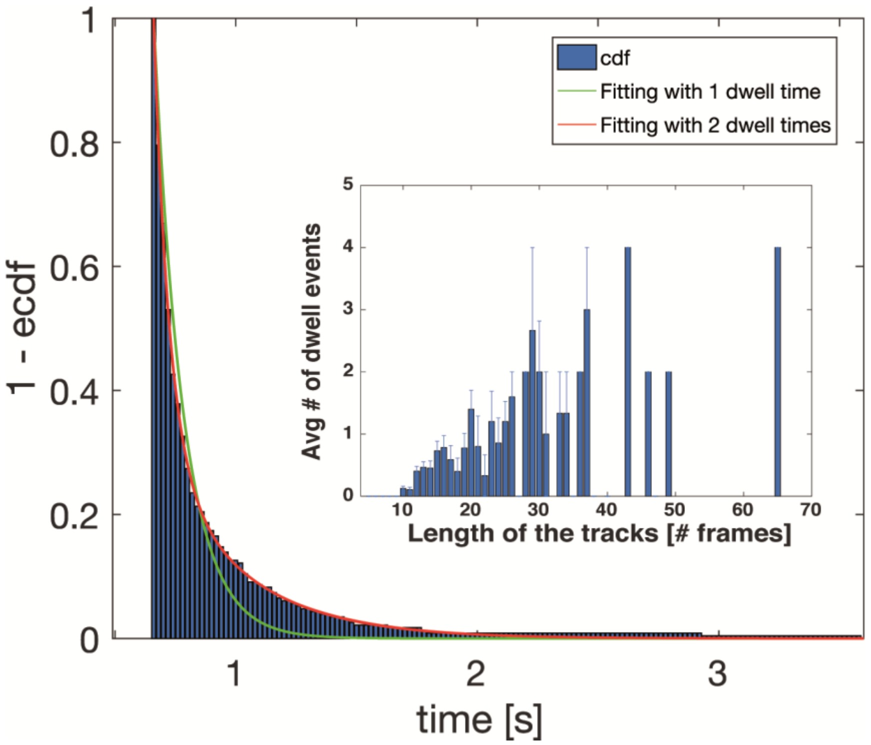

**Fig. S6 Survival function of dwell times for DnaE-mVenus in untreated cells.** Probability that a molecule will remain inside a circle of radius of 120 nm for time t. Dwell times are estimated using an exponential decay model. Blue bars show empirical data, green and red lines show the 1 and 2-component exponential models fitted to the data. A 1-component decay can not explain long dwell times, a better result was obtained for a 2 components model, resulting $\tau_1$= 0.071 s and $\tau_2$= 1.11 s. Inset shows the average number of dwell events per length of the track including standard error.

**TABLE S1** Bacterial Strains and plasmids.

| Strain or Plasmid | Relevant features | Reference or source |
|---|---|---|
| *B. subtilis* | | |
| BG214 | Wild type | |
| PG3300 | amyE::psg1192 *dnaX*-cfp$_{specR}$ | (1) |
| PG3320 | *dnaC*-mVenus$_{cmR}$ | This study |
| PG3302 | *dnaE*-mVenus$_{cmR}$ | (1) |
| PG3321 | *dnaG*-mVenus$_{cmR}$ | This study |
| PG3322 | *dnaX*-cfp$_{specR}$ *dnaC*-mVenus$_{cmR}$ | This study |
| PG3307 | *dnaX*-cfp$_{specR}$ *dnaE*-mVenus$_{cmR}$ | (1) |
| PG3323 | *dnaX*-cfp$_{specR}$ *dnaG*-mVenus$_{cmR}$ | This study |
| *E. coli* | | |
| DH5α | *sup*E44 Δ*lac*U169 ϕ80d*lac*ZΔM15 *hsd*R171 *recA*1 *endA*1 *gyrA*96 *thi*-1 *relA1* | New England Biolabs (NEB) |
| PG3324 | DH5α pSG1164::*dnaC*-mVenus$_{cmR}$ | This study |
| PG3325 | DH5α pSG1164::*dnaG*-mVenus$_{cmR}$ | This study |
| PG3319 | DH5α pSG1192::*dnaX*-cfp$_{specR}$ | (1) |
| PG3315 | DH5α pSG1164::*dnaE*-mVenus$_{cmR}$ | (1) |

**TABLE S2** Oligonucleotides used in this work.

| Name | Sequence[a,b] | Construct |
|---|---|---|
| 22502 | *C*GAATTCATGTCTTTTGTTCACCTGCA | PG3315 |
| 22503 | *A*GGGCCCTTACCACTGTTTTAAAACGA | PG3315 |
| 22604 | *AC*GAATTCGATTTACATCGATGATACAC | PG3324 |
| 22605 | *A*GGGCCCTGCGCCGGGCGGAACGCCTG | PG3324 |
| 22606 | *C*GAATTCGCTGACAATAGCGGTGAAAC | PG3325 |
| 22607 | *A*GGGCCCTTTTAAAGATCGGTTCAATG | PG3325 |

[a] Non-encoded bases introduced as clamps are shown in italics. Restriction sites are underlined; the oligonucleotides carry either *Eco*RI (GAATTC) or *Apa*I (GGGCCC) sites.

[b] The location is indicated by the first 5' nucleotide and the replicon where the sequence is located. Accession numbers are *dnaC* (CP053102 REGION: 4257943…4259307), *dnaE* (CP053102 REGION: 3104280…3107627) and *dnaG* (CP053102 REGION: 2714536…2716347) of *B. subtilis.*

**TABLE S3.** Average dwell times (in seconds).

| | No treated | MMC | HPUra | SHX |
|---|---|---|---|---|
| | | **DnaE-mVenus** | | |
| $\tau_1$ | 0.086±0.05 | 0.088±0.07 | 0.086±0.04 | 0.064±0.03 |
| $\tau_2$ | 1.52±0.12 | 1.4±0.23 | 1.25±0.54 | 1.01±0.44 |
| **Fraction $\tau_1$ (%)** | 23±0.64 | 26±0.83 | 6±0.94 | 3±0.85 |
| **Fraction $\tau_2$ (%)** | 77±0.54 | 74±0.67 | 94±0.73 | 97±0.63 |
| | | **DnaC-mVenus** | | |
| $\tau_1$ | 0.11±0.05 | 0.081±0.09 | 0.082±0.05 | 0.071±0.05 |
| $\tau_2$ | 1.45±0.35 | 1.19±0.48 | 1.20±0.50 | 1.13±0.50 |
| **Fraction $\tau_1$ (%)** | 28±0.45 | 19±0.46 | 15±0.53 | 11±0.51 |
| **Fraction $\tau_2$ (%)** | 72±0.74 | 81±0.63 | 85±0.72 | 89±0.64 |
| | | **DnaG-mVenus** | | |
| $\tau_1$ | 0.071±0.04 | 0.067±0.04 | 0.011±0.05 | 0.075±0.05 |
| $\tau_2$ | 1.11±0.52 | 1.15±0.48 | 1.29±0.56 | 1.11±0.45 |
| **Fraction $\tau_1$ (%)** | 29±0.42 | 19±0.40 | 30±0.38 | 32±0.65 |
| **Fraction $\tau_2$ (%)** | 71±0.62 | 81±0.55 | 70±0.64 | 68±0.56 |

### Supplementary methods

Molecular and microbiological procedures.

Basic DNA manipulations and molecular techniques were done using established procedures (2). To generate specific DNA fragments suitable for cloning, all PCR amplifications were carried out using *Taq* polymerase High Fidelity (New England Biolabs). Amplification protocols consisted of 30 cycles of 1 minute at 94°C, 1 minute at variable temperature (depending on the primer combination), and 1 to 3 minutes at 68°C. The DNAs were resuspended in Tris-EDTA buffer and digested with the appropriate restriction enzyme(s) to generate the required ends of the fragments. The DNA fragments were purified before cloning by isolating them from agarose gels. Primer combinations (Table S2) employed to generate fragments for cloning and electrophoretic mobility shift analyses were generated by annealing custom-made oligonucleotides, purified using a GeneJET extraction kit (QIAGEN). For ligations, T4 polynucleotide ligase (New England Biolabs) was used. Plasmid transformation of *E. coli* was done using $CaCl_2$-competent cells. All plasmid constructions were verified by restriction analysis and PCR and in all of the cases, by DNA sequencing.

Western blotting.

*B. subtilis* cultures (1 ml) were harvested by centrifugation. The pellet was resuspended in lysis buffer (20 mM Tris-HCl [pH 7.0], 10 mM EDTA, 1 mg ml$^{-1}$ lysozyme, 10 g ml$^{-1}$ DNase I, 100 g ml$^{-1}$ RNase I, 1 tablet of Mini EDTA-free, EASY pack (Roche, protease inhibitor cocktail)), and incubated for 30 min at 37°C. Proteins were separated by running 12% sodium dodecyl sulfate-polyacrylamide gel electrophoresis (SDS-PAGE) and were transferred onto nitrocellulose membrane followed by blocking with 5% milk in PBST (80 mM $Na_2HPO_4$, 20 mM $NaH_2PO_4$, 100 mM NaCl, 0.2% (v/v) Tween-20). Proteins were probed using a 1:500 dilution (rabbit-α-GFP) and secondary antibody was added (goat-α-rabbit-antibody in 1:10000 dilution) after a series of washing steps with PBST. Solution A (100 mM Tris pH 8.5, 2.5 mM Luminol, and 0.4 mM Coumaric acid) and Solution B (100 mM Tris pH 8.5, 0.02% (v/v)

$H_2O_2$) were prepared and mixed followed by incubation for 2 min for chemiluminescence detection with ChemiDocTM MP System (BIO-RAD).
